# Oxytocin and shared intentionality drive variation in cooperation in children

**DOI:** 10.1101/2024.02.25.581910

**Authors:** Jennifer McClung, Zegni Triki, Monica Lancheros Pompeyo, Romain Fassier, Yasmin Emery, Adrian Bangerter, Fabrice Clément, Redouan Bshary

## Abstract

While humans cooperate with unrelated individuals to an extent that far outstrips any other species, we also display extreme variation in decisions about whether to cooperate or not. A diversity of cognitive, affective, social, and physiological mechanisms interact to shape these decisions. For example, group membership, shared intentionality talk (i.e. talk about shared goals), and natural initial oxytocin levels affect cooperation in adults in an optimal foraging paradigm that is loosely modelled on the iterated prisoner’s dilemma. In this ‘egg hunt’, shared intentionality talk was key to achieve cooperation, and it occurred more between participants who shared the same group membership and had higher initial oxytocin levels. Such complex interactions raise the question of the age at which humans develop the necessary mechanisms to cooperate effectively in the egg hunt game. Here, we tested children in secondary school aged between 10 and 14 years. We found that, as for adults, shared intentionality talk was crucial for successful cooperation. Furthermore, initial oxytocin levels affected cooperation through shared intentionality talk. In contrast, group membership did not affect behaviour. Finally, pre- and post-experiment oxytocin levels showed various interactions with group membership and gender. Thus, children’s performance was relatively similar to adults while showing some differences with respect to underlying mechanisms. Our study is a rare contribution to further our understanding of the role of oxytocin in early adolescent social behaviour.

## Introduction

The amount and complexity of cooperation humans engage in exceeds that in other species. At the same time, humans are not always cooperative, and between-group competition is a key ecological factor that promotes cooperation within groups, not only in humans but in a variety of other species (De Dreu and Triki, 2022). Strong between-group competition in human history may explain the large literature in social psychology that has documented in-group bias, or favouritism shown to members of one’s own group, across a range of behaviours (Beaupré and Hess, 2003; Lonsdale and North, 2009). Such biases may lead to increased cooperation even within groups that are randomly formed (Charness et al., 2007).

Both psychological and physiological mechanisms may contribute to the formation and expression of ingroup biases. Regarding the former, shared intentionality, defined as the ability to represent mentally another’s goals and then to assume them as one’s own (Tomasello et al., 2005), arises more readily during interactions with ingroup members and thereby increases cooperation (McClung et al., 2017). A key physiological mechanism supporting an in-group bias in cooperation is the evolutionary ancient neuropeptide oxytocin (Triki et al., 2022), which is produced in the hypothalamus in response to social bonding and affiliation (Carter, 2014). Oxytocin has also been co-opted in humans to regulate group living, specifically by facilitating compliance with group norms and increasing trust, empathy, and cooperation between members of the same group (for a review, see (De Dreu and Kret, 2016)).

Mechanisms like shared intentionality and oxytocin production may also impact humans’ cooperative decisions in situations in which to cooperate means to forego immediate benefits, while defect increases the immediate benefits. Such conditions are met in the iterated prisoner’s dilemma (IPD). Game theoreticians, mainly in the field of economics, have spent much effort to determine what kind of conditional cooperative strategies may lead to stable cooperation in the IPD. Meanwhile, psychologists used both modelling approaches and artificial laboratory setups to assess the spontaneous cooperative decisions that humans make in paradigms modelled after real-life situations. In these latter cases, humans may show spontaneous cooperation without considering defection opportunities in games that adhere to an IPD payoff matrix but also allow more natural human social behaviour (e.g. talking and interacting). An example is in the ‘egg hunt’ game (McClung et al., 2018, 2017), a task in which pairs of participants engaged in a 5 min hunt for screws of three different colours (only two of which were rewarded) that were hidden in small, plastic eggs throughout a room. Each participant was rewarded for the total collection of screws of one assigned colour (either red or blue, with green screws being irrelevant to the total reward). Therefore, participants could choose to help the other during the hunt each time they came across a screw of the other’s colour. Crucially, this option was not mentioned in the instructions, and participants, therefore, had to discover spontaneously that the possibility of mutual helping existed. Furthermore, helping was costly (in terms of time taken away from the collection of one’s own screws). However, the egg hunt differs from a typical prisoner’s dilemma game in that it is not framed as a shared game with discrete behavioural options and a 2 x 2 payoff matrix, decisions about what to do with a partner’s screw were not imposed or discussed by experimenters, and they are made individually, and participants can talk in the talking condition.

In the egg hunt, participants did not show any evidence for conditional cooperative strategies (i.e. they did not base their decisions to cooperate on their partner’s cooperation). Instead, talking promoted mutual helping if participants discussed aspects of the egg hunt regarding shared intentionality (meaning shared goals within the hunt), which was more common in ingroup pairs (McClung et al., 2017). In-group and out-group pairs were formed by using a minimal-group paradigm (Tajfel et al., 1971) by giving participants the impression (without overtly lying to them) that their responses to a 10-item questionnaire about food preferences were used to put them into a pair with someone with similar preferences (in-group pair) or a pair with someone with different preferences (out-group pair). In reality, categorisation was random (McClung et al., 2017). In parallel, endogenous baseline oxytocin predicted helping and conversation type, but crucially also as a function of group membership (McClung et al., 2018): higher baseline oxytocin predicted increased helping but only between in-group participants, as well as decreased discussion about individuals’ goals between in-group participants but conversely more of such discussion between out-group participants. The non-talking condition typically yielded no helping, showing that a cooperative solution was much more difficult to achieve in the egg hunt than in a standard IPD game played on computers in a laboratory with all potential decisions explicitly explained by the experimenter.

The complexity of the task, and the mechanisms involved in its solution, make it a more realistic simulacrum of cooperation that nonetheless retains the structure of the IPD. It thus constitutes a useful paradigm to investigate further questions, in particular at what age humans start to develop the capacity to cooperate in this task. Developmental psychology provides many insights into the development of the theory of mind and shared intentionality and its effects on social behaviour. Evidence regarding shared intentionality shows that most of its features are already present at the age of 3 (Tomasello and Gonzalez-Cabrera, 2017), and at the age of about 4 years, children start to pass the litmus test of an explicit theory of mind, the Sally-Anne test (Baron-Cohen et al., 1985; Wimmer and Perner, 1983). Theory of mind correlates with the degree of cooperation children engage in: those who can pass a typical false belief test cooperate more in an ultimatum game (Takagishi et al., 2010), and children who are better at ascribing emotions to others offer more in a dictator game (Gummerum et al., 2010). In contrast to the abundant research on child cognitive development, there is a lack of studies on how hormones and neurohormones affect cooperation in children. To the best of our knowledge, we are unaware of any other study investigating the role of oxytocin on human social behaviour at a young age.

While shared intentionality develops early, Tomasello and Gonzalez-Carebra (2017) emphasise that early use is largely directed at adults, and that usage of shared intentionality in interactions with peers develops later and might still be fine-tuned during the teenage years (13-18 years of age). In line with this assessment, a pilot presentation of the egg hunt game at an Open Day held at the University of Neuchâtel yielded no evidence that children up to the age of 8 years could achieve cooperation in the talking condition. Based on these pilot observations, we decided to test children between the ages of 10 and 14 years of age (adhering to the UN’s definition in the Convention on the Rights of the Child). Participants were all students at a middle school in Neuchâtel, Switzerland, and they carried out the egg hunt in a manner that largely replicated the experimental design of the earlier study (McClung et al., 2018, 2017). Participants were randomly put into talking/non-talking and in-group/out-group conditions in a full factorial design, though rewards differed from the study on adult university students as children received candies rather than money. We were interested in whether we could replicate the previous results found with university students. Thus, we asked to what extent 10-14-year-olds depend on talking (and particularly on shared intentionality talk), initial oxytocin levels, and shared group membership to achieve high levels of cooperation in the egg hunt.

## Methods

### Participants

The University of Neuchâtel’s ethics committee approved the current study (decision letter from 20.09.2016 in supporting online material). Participants were recruited amongst year-seven pupils at Collège du Mail, a secondary school in Neuchâtel, Switzerland. We obtained written consent from children’s parents prior to the experiments. We formed 54 same-sex pairs, N = 108 participants. Due to some participants talking during the non-talking condition, as well as some technical troubles with the eye-tracking equipment, we had to remove pairs (when talking occurred) and individuals (for which the camera did not work, we kept the partner in such cases) from the data set (20 in total). One pair which included a boy and a girl was also discarded from the data. Our final dataset thus contained 86 participants wherein 41 were females and 45 were males, with age range between 10 and 14 years old.

All pairs were tested in single sessions lasting approximately 20 min. Experiments were conducted in French (all methods and electronic supplementary information regarding the procedure are provided as translations into English). All participants were naïve to the experimental hypothesis and were informed that they could stop the experiment at any time. They and their parents were also told that their data would be treated confidentially and used anonymously in publication. We debriefed all participants about the goals of the study directly after their trials. Before debriefing, they had to fill in a questionnaire for us to check how much they had understood about the goals of our study.

### Procedure

The experiment was conducted at the school. Participants were allowed to leave classes for the experiment, and we also ran trials during breaks. We had access to a dedicated room with tables on which we could distribute a total of 100 Kinder^®^ egg shells. Sixty of them contained green screws, which were not rewarded. Twenty eggs contained red screws, and 20 contained blue screws, the collection of which would yield a reward: one candy for each red screw for one participant, one candy for each blue screw for the other participant. Participants were informed of the reward rate before carrying out the experiment.

For our two factorial design with the factors talking condition (talking vs non-talking) and group membership (in-group vs out-group), pairs were randomly assigned to the talking or non-talking condition, and we operationalised group membership by asking participants to complete a short survey with 10 questions about their food preferences, using a 5-point Likert scale from ‘totally disagree’ to ‘totally agree’ (see ‘Food preferences questionnaire’ in electronic supplementary material). The experimenters then made a small show of assessing the surveys and then categorising participants as ‘apple’ or ‘orange’, asking participants to put on a corresponding lab coat to make their group more salient: green with an apple on the pocket for those in the ‘apple’ group, orange with an orange on the pocket for those in the ‘orange’ group. It is important to note that at no point did the experimenters actually discuss the participants’ results on the survey with them or their categorisation (which was in fact random and did not result from the survey results). In-group pairs were formed by categorising both individuals as either ‘orange’ or ‘apple’, while out-group pairs consisted of one ‘orange’ and one ‘apple’. After pairs were formed, participants were fitted with eye-tracking glasses (ETG 2.1 models provided by SensoMotoric Instruments GmbH, Germany) to enable us to record behavioural data.

For the saliva samples, participants had to chew for 2 min on cotton swabs (Salivettesw, Sarstedt, Germany) and then spit the chewed swab into a labelled and sealable vial. We took the first sample to assess baseline oxytocin after participants filled in the food preferences survey and before we categorised them: this was in order to ensure that the group categorisation did not impact baseline oxytocin measures.

After the first saliva sample, we gave participants instructions for the egg hunt (see electronic supplementary material for Instructions and experimental set-up). The instructions informed participants that they would take part in an egg hunt and that each egg had either a red, blue, or green screw within. Participants were given a board into which they could screw the screws they wanted to collect, and they were told that one of them would be rewarded for all the red screws collected (at 1 candy each), that the other would be rewarded for all the blue screws collected (also at 1 candy each), and that the green screws were not rewarded. Participants were told that they could carry out the hunt in any way they wanted. We never explicitly told participants that they could collect the other’s screws or cooperate in any way. Instead, we informed them that their reward would be based on the total number of screws ‘screwed into a board’ without any reference to which board the screws were required to be screwed into. As a final instruction, participants were informed whether they would be allowed to talk freely (‘talking’ condition) or not at all (‘non-talking condition), and they were given a 1 min ‘countdown’ before the hunt started. Again, during this 1 minute, participants in the talking condition were allowed to talk, whereas those in the non-talking condition were not. The egg hunt then lasted 5 minutes after the countdown. Immediately at the end of the hunt, approximately 11 min after the first sample, a second saliva sample was collected using the same method as described above. We used these samples to assess the final oxytocin levels. All saliva samples were frozen at -20°C for hormonal analysis.

### Behavioural data

Following McClung et al. (2017), we quantified the number of events in which individuals provided costly helping, defined as acts that benefit the partner but that take time so that the helper experiences opportunity costs (i.e. carrying the screw to the partner or screwing the other’s screw into one’s own board). In a previous egg hunt experiment, helping at a cost to oneself increased in the in-group condition (McClung et al., 2017), an effect also seen under the artificial external application of oxytocin in another cooperation experiment (De Dreu, 2012).

Conversations in the talking condition were fully transcribed and segmented into units that could be categorised as (i) shared intentionality talk (references to shared goals in the task included both planning and referencing), (ii) individual goal talk (references to separate goals that were not shared), (iii) task talk (practical aspects without any link to goals), or (iv) other (all utterances that could not be classified in one of the other three categories) (McClung et al., 2017).

### Hormonal data

Oxytocin analyses followed procedures by McClung et al. (2018), using similar infrastructure and equipment at the University of Neuchâtel. Briefly, we first ran the collected saliva, from Salivettes®, through extraction columns WCX 25mg 1 mL columns (Evolute SPE, nr. 602-0002-A) to allow for capturing the oxytocin molecules. Then, the extracts were assessed with the enzyme immunoassay technique ELISA to detect the levels of oxytocin in pg per 1 mL of saliva (for further information, see detailed protocol in the Supplementary Material).

### Data analyses

For statistical analyses, we estimated the proportion of helping as the number of screws transferred to the partner from the total screws found of the partner’s colour. Following McClung et al. (2018), the proportions were converged to a binary outcome, that is, proportions < 0.5 were scored as zero helping, whereas proportions ≥ 0.5 were scored as total helping or “1”. As for the types of conversation, we retained for the statistical analyses three types of talk in the form of proportions from the total talk: shared intentionality talk, individual goal talk, and task talk.

The statistical software R version 4.2.1 (R Core Team, 2022) was used to generate the statistics and figures. Before proceeding with data analyses linked to our hypotheses testing, we explored whether potential confounding covariation between baseline oxytocin levels and either age of the participants, gender, group membership or talking condition may have existed in the data (see supplementary material and supplementary Figure S1). For statistical models, we ran a set of Bayesian generalised linear mixed-effects models (BGLMM), and Markov chain Monte Carlo sampler for multivariate generalised linear mixed models (MCMCGLMM). Throughout all tests, pairs’ identity was fitted as the random factor. *Post hoc* analyses were performed with the emmeans package in R wherever appropriate.

Given the multiple testing of “helping behaviour” (i.e., in five statistical models) and “change in oxytocin levels” (i.e., in four statistical models), we employed the False Discovery Rate (FDR) approach to set a significance threshold adapted to every *p*-value (Benjamini and Hochberg, 1995). To do so, we set a derived significance threshold for *p*-values by using the following function: (*i/m*)*Q*; with *i*: *p*-value rank, *m*: number of comparisons (five for helping behaviour, and four for change in oxytocin levels), and *Q*: maximum acceptable FDR set at alpha = 0.05. For further details, step-by-step statistical code is archived along with data at the public repository Figshare (see Data and Code accessibility statement).

## Results

### Helping behaviour

We tested whether helping behaviour differed between participants as a function of our two-factorial design: talking condition (talking vs non-talking) and group membership (out-vs in-group). There was a significant main effect of talking where participants in the talking condition had higher helping scores than those in the non-talking condition (BGLMM, N = 86, estimate [lower and upper 95 % confidence interval limits] = 7.660 [2.75, 12.60], *p* = 0.002, Fig. 1a). This effect was independent of group membership where being in-group or out-group participant did not significantly affect levels of helping (BGLMM, N = 86, 1.026 [-3.01, 5.07], *p* = 0.618, Table 1, Fig. 1).

**Table 1.** Summary table of the main results for helping behaviour. Values indicated in bold are statistically significant (*p* ≤ *FDR derived significance threshold)*.

| Model class | term | N | Estimate /<br>posterior<br>mean* | Lower limit<br>95% CI | Upper limit<br>95% CI | p-value | p-value<br>rank (i) | FDR derived<br>significance<br>threshold | Explained<br>variance (<br>Marginal,<br>conditional<br>R<sup>2</sup>; or<br>pseudo R<sup>2</sup>*) |
| --- | --- | --- | --- | --- | --- | --- | --- | --- | --- |
| Bayesian GLMM | (Intercept) | 86 | **-4.260** | **-7.80** | **-0.73** | **0.018** | **5** | **0.042** | **0.42, 0.80** |
|  | group (out-group) |  | 1.030 | -3.01 | 5.07 | 0.619 | 33 | 0.275 |  |
|  | **talking (talk)** |  | **7.660** | **2.75** | **12.60** | **0.002** | **1** | **0.008** |  |
|  | gender (male) |  | -2.060 | -7.37 | 3.25 | 0.448 | 27 | 0.225 |  |
|  | age |  | -0.149 | -1.21 | 0.91 | 0.783 | 37 | 0.308 |  |
|  | group (out) x talking (talk) |  | -4.680 | -9.93 | 0.57 | 0.081 | 10 | 0.083 |  |
|  | group (out) x gender (male) |  | 2.790 | -2.22 | 7.80 | 0.275 | 20 | 0.167 |  |
|  | talking (talk) x gender (male) |  | -1.910 | -6.85 | 3.03 | 0.448 | 28 | 0.233 |  |
| Bayesian GLMM | (Intercept) | 78 | -9.670 | -21.50 | 2.13 | 0.108 | 13 | 0.108 | **0.76, 0.95** |
|  | baseline oxytocin |  | -4.910 | -10.20 | 0.38 | 0.069 | 8 | 0.067 |  |
|  | group (out-group) |  | 7.120 | -4.82 | 19.10 | 0.242 | 19 | 0.158 |  |
|  | talking (talk) |  | 13.600 | 0.44 | 26.70 | 0.043 | 7 | 0.058 |  |
|  | gender (male) |  | -6.620 | -16.60 | 3.33 | 0.192 | 16 | 0.133 |  |
|  | baseline oxytocin x group (out) |  | 3.100 | -1.32 | 7.53 | 0.169 | 15 | 0.125 |  |
|  | **baseline oxytocin x talking (talk)** |  | **4.900** | **0.26** | **9.54** | **0.039** | **6** | **0.050** |  |
|  | baseline oxytocin x gender (male) |  | -0.797 | -4.37 | 2.77 | 0.662 | 34 | 0.283 |  |
|  | group (out) x talking (talk) |  | -11.700 | -25.10 | 1.70 | 0.087 | 11 | 0.092 |  |
|  | group (out) x gender (male) |  | 4.730 | -3.09 | 12.60 | 0.236 | 18 | 0.150 |  |
|  | talking (talk) x gender (male) |  | 1.220 | -7.07 | 9.51 | 0.773 | 36 | 0.300 |  |
| **MCMCglmm\*** | (Intercept) | 37 | 16.534 | -4.66 | 43.04 | 0.141 | 14 | 0.117 | **0.95** |
|  | **shared intentionality** |  | **44.324** | **2.64** | **90.96** | **0.005** | **2** | **0.017** |  |
|  | group (out) |  | -8.270 | -31.42 | 11.16 | 0.383 | 24 | 0.200 |  |
|  | gender (male) |  | -6.781 | -33.41 | 20.14 | 0.545 | 30 | 0.250 |  |
|  | **shared intentionality x group (out)** |  | **35.582** | **5.93** | **75.99** | **0.017** | **4** | **0.033** |  |
|  | shared intentionality x gender (male) |  | -23.959 | -78.82 | 21.15 | 0.314 | 22 | 0.183 |  |
| **MCMCglmm** | (Intercept) | 37 | 12.684 | -31.35 | 68.37 | 0.712 | 35 | 0.292 | 0.39 |
|  | **individual goal talk** |  | **-54.435** | **-113.85** | **-6.05** | **0.007** | **3** | **0.025** |  |
|  | group (out) |  | -27.995 | -112.12 | 24.09 | 0.591 | 32 | 0.267 |  |
|  | gender (male) |  | -15.234 | -71.36 | 28.37 | 0.568 | 31 | 0.258 |  |
|  | individual goal talk x group (out) |  | 30.518 | -1.99 | 67.41 | 0.080 | 9 | 0.075 |  |
|  | individual goal talk x gender (male) |  | 31.749 | -38.76 | 114.93 | 0.388 | 25 | 0.208 |  |
| **MCMCglmm** | (Intercept) | 37 | 28.676 | -29.04 | 105.40 | 0.324 | 23 | 0.192 | 0.30 |
|  | task talk |  | -22.439 | -69.89 | 27.57 | 0.497 | 29 | 0.242 |  |
|  | group (out) |  | -24.939 | -101.33 | 41.67 | 0.438 | 26 | 0.217 |  |
|  | gender (male) |  | -37.151 | -111.12 | 31.38 | 0.233 | 17 | 0.142 |  |
|  | task talk x group (out) |  | -33.519 | -97.02 | 21.96 | 0.292 | 21 | 0.175 |  |
|  | task talk x gender (male) |  | 54.109 | -6.49 | 119.24 | 0.092 | 12 | 0.100 |  |

*\*MCMCGLMM models helped in fitting data with perfect separation. The estimated marginal- $R^2$ is specific to the model. For MCMGLMM we report the posterior mean instead of estimate.*

**Figure 1.**
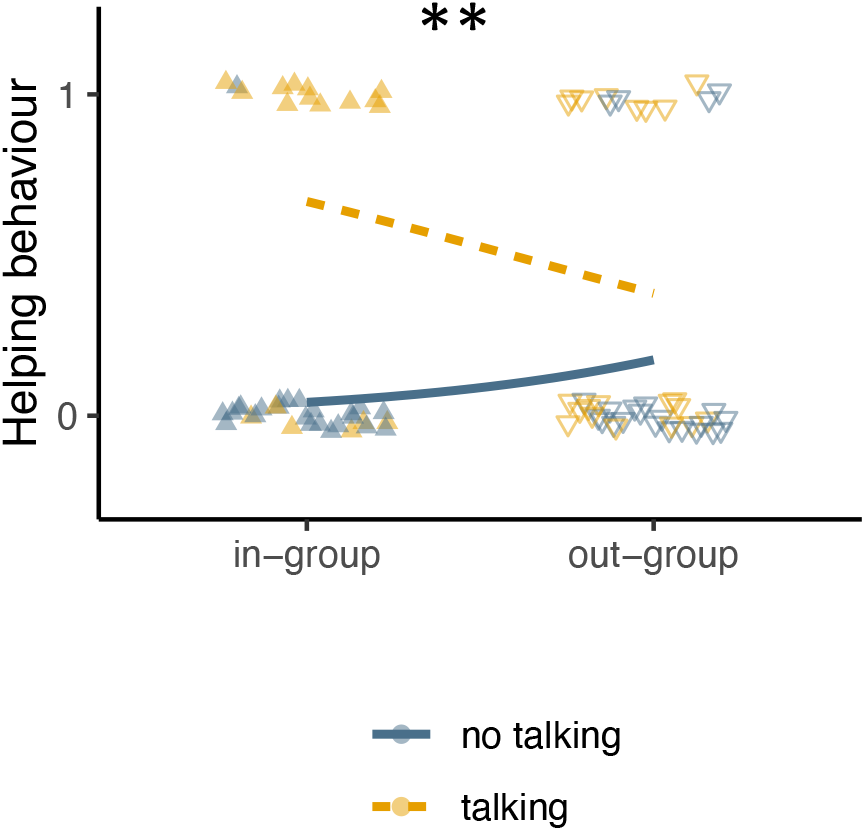
Helping behaviour as a function of group membership and talking. Logistic regression lines of helping behaviour (binomial variable with 0 = mostly no helping and 1 = mostly helping) for participants in the “talking” vs “non-talking” condition. BGLMM ^**^*p* < 0.01.

### Effects of oxytocin on helping behaviour

There was no significant difference in baseline oxytocin levels as a function of gender, age, group membership, or talking condition of the participants (see supplementary material and supplementary Figure S1).

Assessing whether baseline oxytocin levels predicted participants’ helping behaviour showed an apparent significant relationship only in interaction effect with the talking condition (BGLMM, N = 78, 4.898 [0.259 9.54], *p* = 0.038, Table 1, Fig. 2a). *Post hoc* analyses showed that, in the non-talking condition, helping behaviour decreased with the increase in baseline oxytocin levels (emtrend = -3.76 [-7.51, - 0.01], *p* = 0.049). In the talking condition, although there was an increase in the helping behaviour when baseline oxytocin increased, the effect was not statistically significant (emtrend = 1.14 [-1.21, 3.48], *p* = 0.342). Baseline oxytocin levels did not significantly affect levels of helping behaviour alone or as a function of group membership (detailed statistics in Table 1). Furthermore, change in oxytocin levels throughout the experiment was not significantly affected by helping behaviour, talk condition and group membership (detailed statistics in Table 2, Fig. 2b).

**Table 2.**
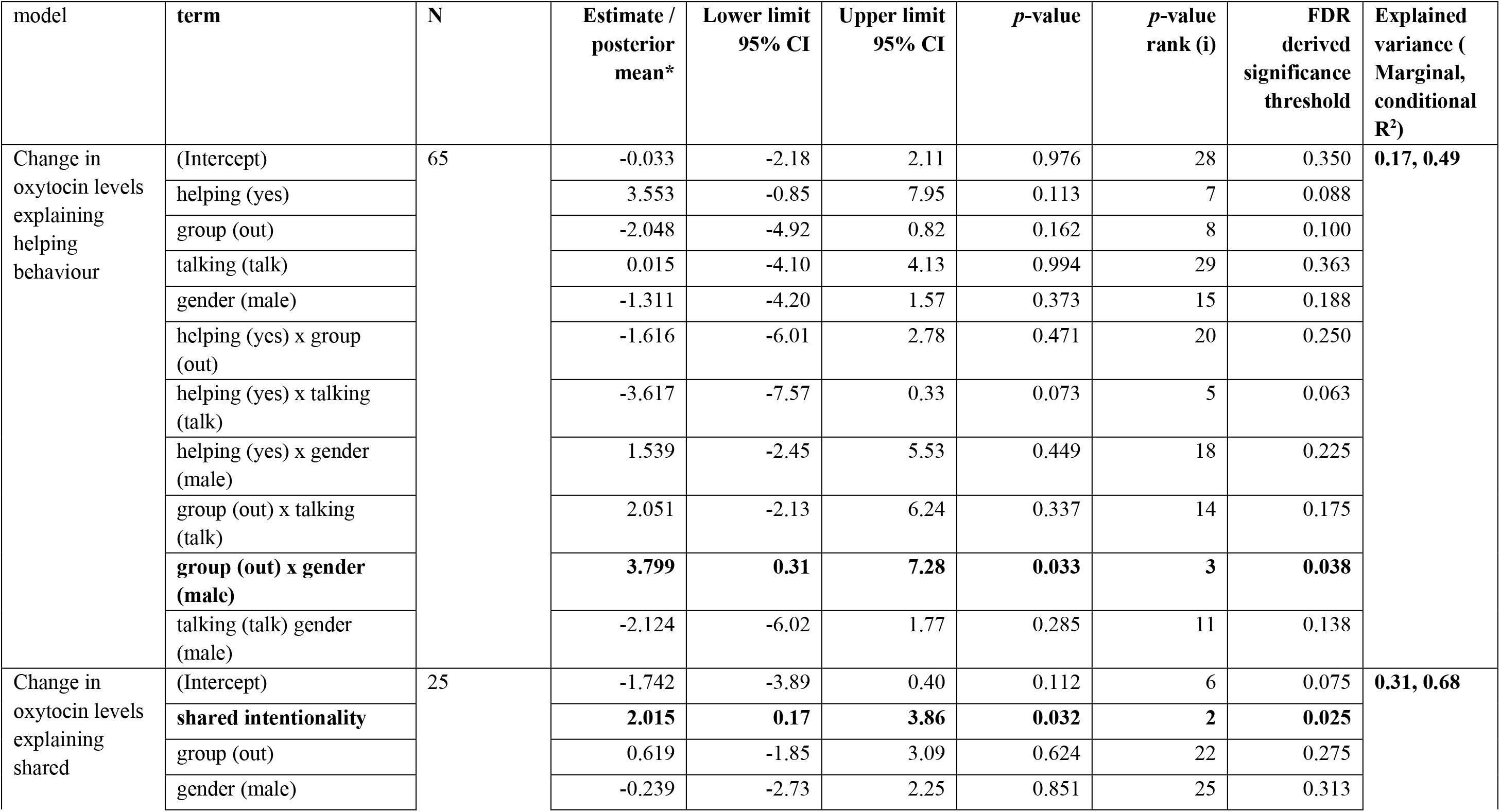
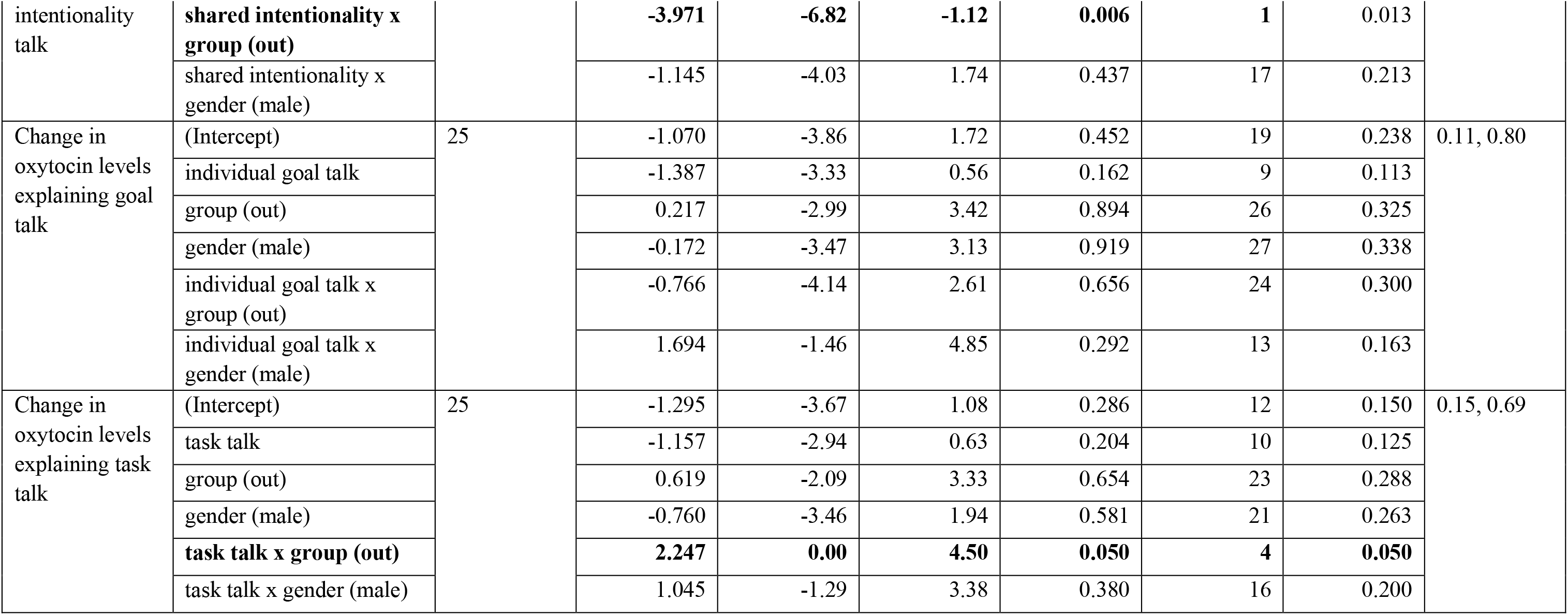
Summary table of the main results for change in oxytocin levels with Bayesian GLMM models. Values indicated in bold are statistically significant (*p* ≤ *FDR derived significance threshold)*.

**Figure 2.**
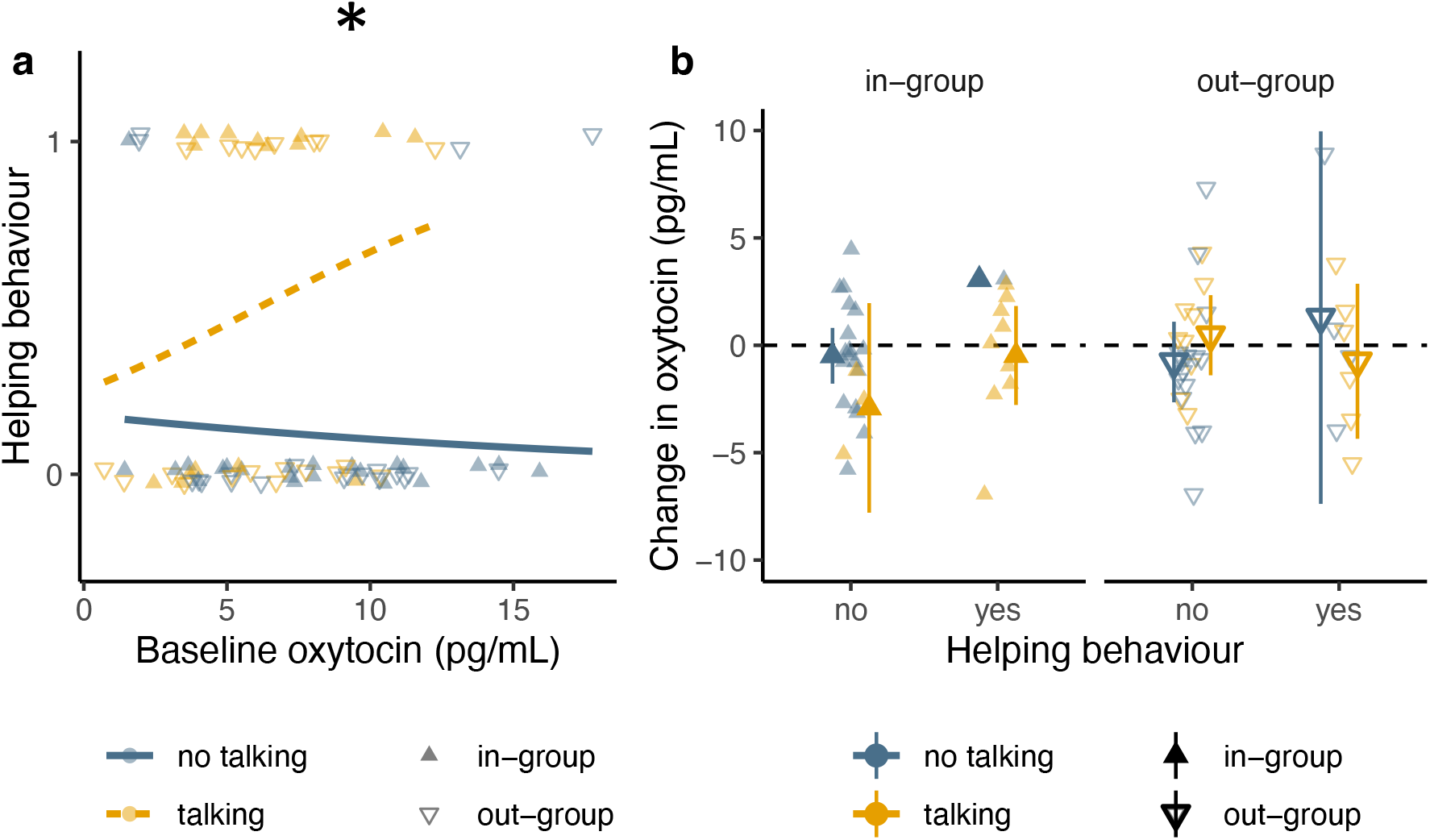
Oxytocin as a mechanism for helping behaviour. (**a**) Relationship of baseline oxytocin to helping behaviour as a function of talking condition (logistic regression lines shown for each talk condition). (**b**) Helping behaviour predicts changes in oxytocin levels (mean and 95% CI) as a function of talking condition and group membership. BGLMM ^*^*p* < 0.05.

### Effects of types of conversation on helping behaviour

For participants in the talking condition, we tested whether their helping behaviour could be predicted by types of talk: shared intentionality, individual goal, and task talk. We found a significant positive effect of shared intentionality talk on helping behaviour (MCMCglmm, N = 37, 44.324 [2.64, 90.96], *p* = 0.005, Fig. 3a). This effect was statistically influenced by group membership (MCMCglmm, N = 37, 35.582 [5.93, 75.99], *p* = 0.017, Fig. 3a). *Post hoc* analyses showed that although shared intentionality facilitated helping behaviour in both out- and in-group participants, the effect was stronger in the out-group (emtrends: in-group, 34.1 [3.59, 58.4]; out-group, 67.2 [21.43, 119.0], Fig. 3a). The statistical analyses of the other two types of talk revealed that individual goal talk was negatively associated with helping levels (MCMCglmm, N = 37, -54.435 [-113.85, -6.05], *p* = 0.007, Fig. 3b) with no effect of task talk (MCMCglmm, N = 37, -22.439 [-69.89, 27.57], *p* = 0.497, Fig. 3c) (see Table 1 for further statistics).

**Figure 3.**
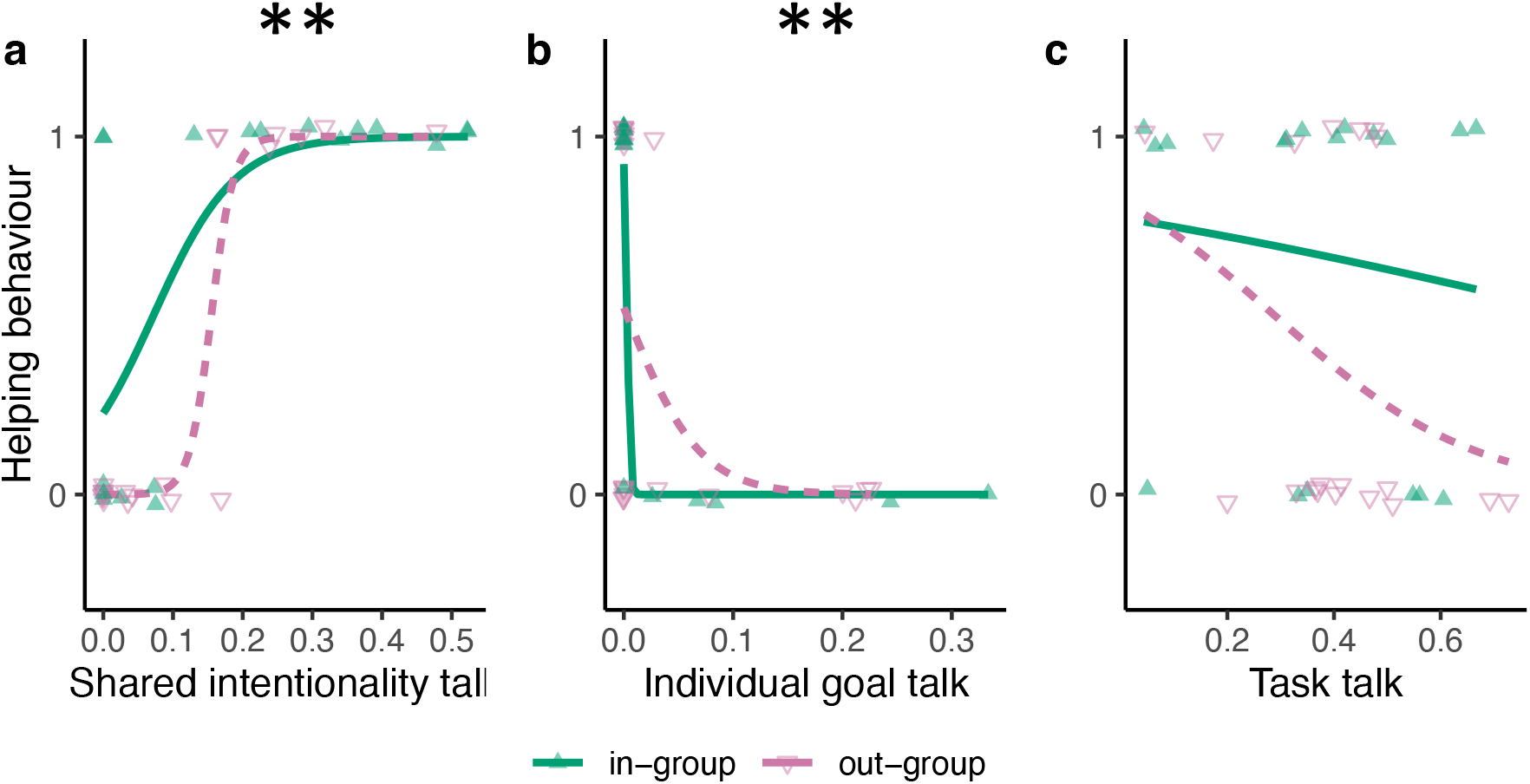
Types of conversation as a mechanism for helping behaviour. Logistic regression lines of helping behaviour as a function of group membership and the three types of talk: (**a**) shared intentionality talk, (**b**) individual goal talk, and (**c**) task talk. MCMglmm ^**^*p* < 0.01.

### The relationship between types of conversation and oxytocin

We explored whether baseline oxytocin levels had any effect on types of talk. We found no significant relationship between baseline oxytocin and types of talk (Fig. 4a-c), we report the detailed statistics in the Supplementary Table S1.

**Figure 4.**
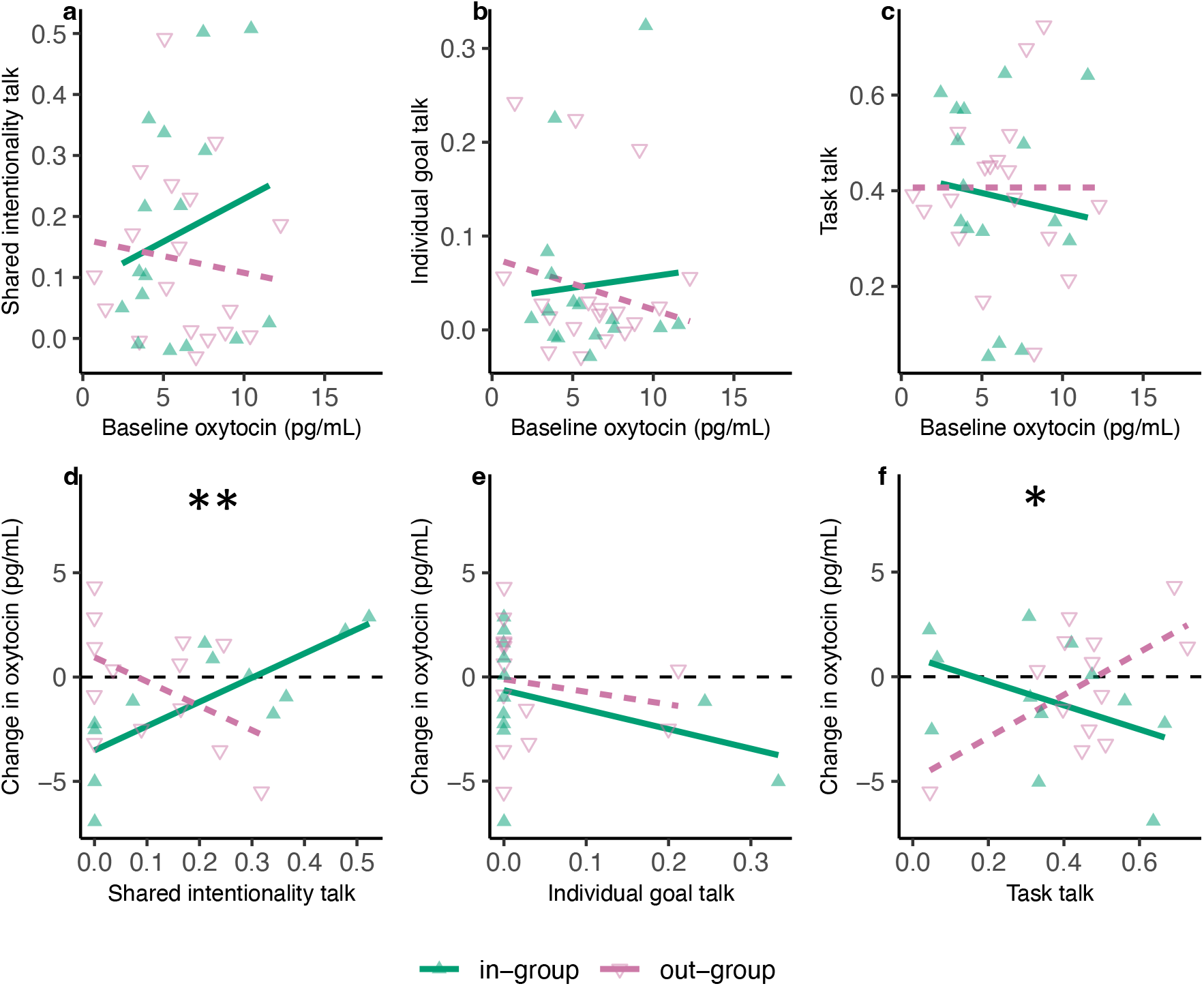
The relationship between types of conversation and oxytocin. Baseline oxytocin a function of group membership predicting the three types of talk: (**a**) shared intentionality talk, (**b**) individual goal talk, and (**c**) task talk. The three types of talk predicting changes in oxytocin levels as a function of group membership are (**d**) shared intentionality talk, (**e**) individual goal talk, and (**f**) task talk. BGLMM ^*^*p* < 0.05, ^**^*p* < 0.01.

Regarding whether types of talk had any effect on change in oxytocin levels throughout experiment, we found that overall shared intentionality talk was positively associated with increased oxytocin levels (BGLMM, N = 25, 2.015 [0.17, 3.86], *p* = 0.032, Fig. 4d). Nevertheless, this effect was group-dependent (BGLMM, N = 25, -3.971 [-1.12, -6.82], *p* = 0.006). The overall significant interaction was driven by two opposing non-significant slopes: in-group participants who used more shared intentionality talk showed a greater increase in oxytocin-from baseline levels (emtrend = 1.44 [-0.45, 3.33], *p* = 0.123) while out-group participants who used more shared intentionality showed a greater decrease in oxytocin from baseline levels (emtrend = -2.53 [-5.29, 0.23], *p* = 0.069) (Fig. 4d). We found an opposite interaction effect for task talk on OT levels (BGLMM, N = 25, 2.247 [0.00, 4.50], *p* = 0.05, Fig. 4f), against based on two non-significant slopes: in-group participants who used more task talking showing a greater decrease in oxytocin from baseline levels (emtrend = -0.635 [-2.3, 1.04], *p* = 0.436) while out-group participants who used more task talk showing a greater increase in oxytocin from baseline levels (emtrend = 1.612 [-4.28, 3.65], *p* = 0.114). There were no significant effects detected for individual goal talk (Table 2, Fig. 4e).

### 4. Gender effect

Throughout all the statical analyses, participants’ gender (male vs female) did not affect helping behaviour (Table 1), or types of talk except for individual goal talk where males showed more of this type of conversation than females (supplementary Table S1). The most evident gender effect was on changes in oxytocin levels as a function of group membership (BGLMM, N = 65, 3.799 [0.31, 7.28], *p* = 0.033, Table 2). Female and male participants had opposite directions of oxytocin changes. Females in the in-group condition had increased oxytocin levels, while males had a decrease, but for the out-group participants, the reverse was true (Supplementary Figure S2).

## Discussion

We investigated the degree to which children aged 10-14 show the capacity to cooperate in the egg-hunt game, a game in which players must solve an iterated prisoner’s dilemma to achieve cooperation, but without being made aware of the payoff matrix. Based on prior positive results, we investigated how group membership, talking, and endogenous oxytocin levels affect levels of cooperation. Furthermore, we investigated which type of talking enhanced cooperation, and finally how our variables of interest would affect change in oxytocin levels. Most analyses can be compared to previously published results on adult participants collected in the same city (McClung et al., 2018, 2017).

As for adults, we found that the possibility to talk had a large positive effect on cooperation and that this was achieved specifically through shared intentionality talk. That is, to cooperate, children too need to both perceive another’s goal as a goal they could share in, and then overtly discuss these shared goals with their partner. Furthermore, the factor talking also interacted with baseline oxytocin levels to increase cooperation (not tested in adults): those children who could talk to each other tended towards more cooperation with increasing baseline oxytocin, whereas amongst children who could not talk, cooperation decreased with increasing baseline oxytocin levels.

The main difference between children and adults was that the children’s cooperation was not affected by their group membership: that is, being in the in-group with a partner did not increase cooperation amongst children the way it did in adults. Furthermore, the experiment affected the children’s change in oxytocin levels in various ways that were absent in adults. All significant results that we discovered for children, for adults or for both are presented in Supplementary Table S2.

We based the choice of the participant’s age range (10-14 years) on previous pilot observations that children up to 8 years old do not achieve any cooperation in the task. As our results were not affected by the children’s age, we tentatively conclude that, at least in Swiss (Western) society, some important improvement occurs with respect to shared intentionality application in later childhood (see age classes in (Tomasello and Gonzalez-Cabrera, 2017)), i.e., between 8 and 10 years of age. One reported change at this age range is that only children older than 8 years can verbally explain the self-reputational consequences of various rule violations (Banerjee et al., 2010), though already 5-year-olds show evidence for behavioural reputation management (Engelmann et al., 2012). Similarly, although younger children may understand aspects of joint commitment (Gräfenhain et al., 2009) and how to choose the most cooperative partner (Grueneisen et al., 2023), the ability to understand promises, how to form commitments and the consequences they entail manifests only at 9 to 10 years of age (Maas and Abbeduto, 2001; Mant and Perner, 1988). Thus, it appears that the advanced use of language, in this case, to express shared intentionality, could be the key cognitive mechanism promoting successful cooperation in the egg hunt from 10 years onwards. Indeed, shared intentionality talk was crucial to achieve cooperation in the task.

Interestingly, shared group membership did not increase levels of cooperation in children as it did in adults (McClung et al., 2017). We obtained these differences despite using the same mechanism of forming in-group and out-group pairs, a minimal group paradigm seemingly based on food preferences but which was in reality arbitrary (McClung et al., 2017). Potentially, the children knew each other, at least by sight, as they were all from the Neuchâtel region around the school, whereas the adults tested may have come from other regions of Switzerland or other countries. Alternatively, the children’s shared identity as students may have been enhanced simply as a result of being in an unfamiliar situation with an unknown experimenter present. It is possible that such strengthening of the children’s shared identity could have dominated our attempts to impose another identity on them. As many studies report in-group favouritism in children from even younger ages (e.g. age 6 (Bennett et al., 2004)), it would appear that our categorisation method did not work as it did with adults and that the negative results on the impact group membership in children’s cooperation should be treated with caution.

Baseline oxytocin levels affected levels of cooperation in interaction with talking, which goes contrary to the reported effects in adults: in the egg hunt with adults, baseline oxytocin interacted with group membership to impact cooperation (McClung et al., 2018). This result has also been obtained in other studies on adult humans where the nasal application of OT promoted ingroup cooperation (De Dreu et al., 2011, 2010). The comparison between children and adults may be partly confounded by the fact that the study on adults could not test for an interaction between oxytocin and talking because of sample size issues. Importantly though, our results show that oxytocin can affect cooperative behaviour in 10-14-year-old children. Interestingly, group membership interacted with three other variables –gender, shared intentionality talk, and task talk – to affect change in oxytocin levels in children. At this stage, we find it difficult to interpret these results. The main possible conclusion is that oxytocin levels are modulated by social conditions and communication content in old children/young teenagers. There is research on oxytocin and conduct problems (Andreou et al., 2018) and mental illness in teenagers (Netherton and Schatte, 2011), but to our knowledge, no other studies to date have examined the effects of oxytocin on the social behaviour of ‘normally’ developing children.

In conclusion, we provide a rare study on cooperation in 10-14-year-old children and the underlying linguistic and physiological mechanisms. The children relied on shared intentionality talk to achieve stable cooperation, similar to what has been documented in adults performing this task. Furthermore, the data clearly show that the children’s oxytocin levels affect and are affected by actions and language use in the experiment. The lack of effect of group membership on cooperation should be investigated in future studies. Furthermore, children ranging from 7 – 10 years of age should be studied systematically to determine at what age the ability to cooperate based on shared intentionality talk develops.

## Ethics statement

The study was approved by the ethics committee at the University of Neuchatel.

## Author contribution

R.B., J.S.M., Z.T., A.B. and F.C. designed the study. J.S.M., M.L.P and R.F. collected behavioural data. Z.T. and Y.E. ran the hormonal analyses. Z.T. analysed the data and generated the figures. R.B. and Z.T. wrote the manuscript with input from all authors.

## Acknowledgement

Funding was provided by the Swiss National Science Foundation (grant no. 31003A_153067/1 to R.B.).

## References

Andreou, D., Comasco, E., Åslund, C., Nilsson, K.W., Hodgins, S., 2018. Maltreatment, the oxytocin receptor gene, and conduct problems among male and female teenagers. Front. Hum. Neurosci. 12, 112.

Banerjee, R., Bennett, M., Luke, N., 2010. Children’s reasoning about the self‐presentational consequences of apologies and excuses following rule violations. Br. J. Dev. Psychol. 28, 799–815.

Baron-Cohen, S., Leslie, A.M., Frith, U., 1985. Does the autistic child have a “theory of mind”? Cognition 21, 37–46.

Beaupré, M.G., Hess, U., 2003. In my mind, we all smile: A case of in-group favoritism. J. Exp. Soc. Psychol. 39, 371–377.

Benjamini, Y., Hochberg, Y., 1995. Controlling the false discovery rate: a practical and powerful approach to multiple testing. J. R. Stat. Soc. Ser. B Methodol. 57, 289–300.

Bennett, M., Barrett, M., Karakozov, R., Kipiani, G., Lyons, E., Pavlenko, V., Riazanova, T., 2004. Young children’s evaluations of the ingroup and of outgroups: A multi‐national study. Soc. Dev. 13, 124–141.

Carter, C.S., 2014. Oxytocin Pathways and the Evolution of Human Behavior. Annu. Rev. Psychol. 65, 17–39. 10.1146/annurev-psych-010213-115110

Charness, G., Rigotti, L., Rustichini, A., 2007. Individual behavior and group membership. Am. Econ. Rev. 97, 1340–1352.

De Dreu, C.K.W., 2012. Oxytocin modulates cooperation within and competition between groups: An integrative review and research agenda. Horm. Behav. 61, 419–428. 10.1016/j.yhbeh.2011.12.009

De Dreu, C.K.W., Greer, L.L., Handgraaf, M.J.J., Shalvi, S., Van Kleef, G.A., Baas, M., Ten Velden, F.S., Van Dijk, E., Feith, S.W.W., 2010. The Neuropeptide Oxytocin Regulates Parochial Altruism in Intergroup Conflict Among Humans. Science 328, 1408–1411. 10.1126/science.1189047

De Dreu, C.K.W., Greer, L.L., Van Kleef, G.A., Shalvi, S., Handgraaf, M.J.J., 2011. Oxytocin promotes human ethnocentrism. Proc. Natl. Acad. Sci. 108, 1262–1266. 10.1073/pnas.1015316108

De Dreu, C.K.W., Kret, M.E., 2016. Oxytocin Conditions Intergroup Relations Through Upregulated In-Group Empathy, Cooperation, Conformity, and Defense. Biol. Psychiatry 79, 165–173. 10.1016/j.biopsych.2015.03.020

De Dreu, C.K.W., Triki, Z., 2022. Intergroup conflict: origins, dynamics and consequences across taxa. Philos. Trans. R. Soc. B Biol. Sci. 377, 20210134. 10.1098/rstb.2021.0134

Engelmann, J.M., Herrmann, E., Tomasello, M., 2012. Five-year olds, but not chimpanzees, attempt to manage their reputations. PLoS One 7, e48433.

Gräfenhain, M., Behne, T., Carpenter, M., Tomasello, M., 2009. Young children’s understanding of joint commitments. Dev. Psychol. 45, 1430.

Grueneisen, S., Török, G., Wathiyage Don, A., Ruggeri, A., 2023. Young children’s adaptive partner choice in cooperation and competition contexts. Child Dev.

Gummerum, M., Hanoch, Y., Keller, M., Parsons, K., Hummel, A., 2010. Preschoolers’ allocations in the dictator game: The role of moral emotions. J. Econ. Psychol. 31, 25–34.

Lonsdale, A.J., North, A.C., 2009. Musical taste and ingroup favouritism. Group Process. Intergroup Relat. 12, 319–327.

Maas, F.K., Abbeduto, L., 2001. Children’s judgements about intentionally and unintentionally broken promises. J. Child Lang. 28, 517–529.

Mant, C.M., Perner, J., 1988. The child’s understanding of commitment. Dev. Psychol. 24, 343.

McClung, J.S., PlacÍ, S., Bangerter, A., Clément, F., Bshary, R., 2017. The language of cooperation: shared intentionality drives variation in helping as a function of group membership. Proc. R. Soc. B Biol. Sci. 284, 20171682. 10.1098/rspb.2017.1682

McClung, J.S., Triki, Z., Clément, F., Bangerter, A., Bshary, R., 2018. Endogenous oxytocin predicts helping and conversation as a function of group membership. Proc. R. Soc. B Biol. Sci. 285, 20180939. 10.1098/rspb.2018.0939

Netherton, E., Schatte, D., 2011. Potential for oxytocin use in children and adolescents with mental illness. Hum. Psychopharmacol. Clin. Exp. 26, 271–281.

R Core Team, 2022. A language and environment for statistical computing.

Tajfel, H., Billig, M.G., Bundy, R.P., Flament, C., 1971. Social categorization and intergroup behaviour. Eur. J. Soc. Psychol. 1, 149–178.

Takagishi, H., Kameshima, S., Schug, J., Koizumi, M., Yamagishi, T., 2010. Theory of mind enhances preference for fairness. J. Exp. Child Psychol. 105, 130–137.

Tomasello, M., Carpenter, M., Call, J., Behne, T., Moll, H., 2005. Understanding and sharing intentions: The origins of cultural cognition. Behav. Brain Sci. 28, 675–691.

Tomasello, M., Gonzalez-Cabrera, I., 2017. The Role of Ontogeny in the Evolution of Human Cooperation. Hum. Nat. 28, 274–288. 10.1007/s12110-017-9291-1

Triki, Z., Daughters, K., De Dreu, C.K.W., 2022. Oxytocin has ‘tend-and-defend’ functionality in group conflict across social vertebrates. Philos. Trans. R. Soc. B Biol. Sci. 377, 20210137. 10.1098/rstb.2021.0137

Wimmer, H., Perner, J., 1983. Beliefs about beliefs: Representation and constraining function of wrong beliefs in young children’s understanding of deception. Cognition 13, 103–128.

