## Supplementary Material for "Oxytocin and shared intentionality drive variation in cooperation in children"

**Supplementary Methods and Material:**

**Hormonal Analyses Procedure:**

<u>Salivary Oxytocin Analysis</u>

The Salivettes® absorb non stimulated saliva passively in the participant's mouth in order to have a minimum of 1mL of saliva. Samples were stored immediately on ice until the end of each trial. Salivary oxytocin was analyzed in the lab at the University of Neuchâtel using the enzyme immunoassay technique ELISA. Frozen samples at -20°C were thawed in 4°C. We tried to handle the sample for most of the following steps in 4°C to insure the stability of the salivary oxytocin. The Salivette® tubes were centrifuged for 10 min at 4°C in 5000 rpm. 1 mL of saliva was mixed with 100 µL of phosphoric acid, then centrifuged at 4°C for 10 min at 17000 rpm to avoid pipetting precipitates. At this step, samples can be stored again at -80 °C for further analyses or processed directly for extraction. For ELISA results to be of any use in detecting oxytocin as opposed to other substances in our samples, extraction is crucial. We therefore conducted extraction with an improved extraction protocol employing WCX 25mg 1 mL columns (Evolute SPE, nr. 602-0002-A) where the protocol was tested on pilot samples prior to the experimental samples. In order to obtain the oxytocin levels in the extracted saliva, we used the Enzo Life Sciences® Oxytocin ELISA kit. Kit preparation steps were from the protocol provided with the kit. To improve the results, we used a BioTek EL x 50 microplate washer. For the plate reading we used a BioTek SynergyHT microplate reader. Finally, to analyze the readings we used the software KC4 (V 3.0). Samples were run in duplicate horizontally. The oxytocin concentrations were then estimated from readings of optical density of the standardized curve with serial dilutions of known concentrations. Following instructions provided by the manual in the kit, optical density readings with coefficients of variation (CV) higher than 15 % were excluded from the data set. Furthermore, samples that fell below the detection range were reported as the lowest level of the sensitivity of the kit, 15 pg mL-1 (personal communication with the technical support of Enzo®) only when the lowest limit detection of the plate is equal to or above the sensitivity of the kit. In total 22 samples fell below the lowest range and were recorded as not available data. Finally, all oxytocin concentrations were corrected by the concentration factor during extraction. The range of CV% of all the oxytocin readings was between 0 and 7.34%. The lowest detectable level was 0.15 pg mL-1, the intra- and inter-assay coefficient of variations were 1.7 % and 0.14 % respectively.

<u>Solid phase extraction</u>

We collected 0.3-2mL of saliva using Sarsted Salivettes®. Salivettes® tubes were centrifuged at 5000 rpm at 4°C for 10 minutes or more. Then, we mix each 1 ml of the saliva with 100 ul of 0.5 N phosphoric acid in Ependorf® tubes 2 ml or 1.5 ml. Ependorf® tubes were centrifuged in the microcentrifuger at 4°C, 17000 rpm for 10 min and the supernatant was recovered in new Ependorf® tubes. Samples were stored at -80°C for further analyses or processed immediately.

We used Biotage® Pressure +48, positive pressure manifold for solid phase extraction employing mixed mode of non-polar/weak cation exchange columns (Evolute SPE biotage 25mg WCX 1ml, nr. 602-0002-A). At this point we tested the C18 columns as described in the kit protocol but there were no oxytocin detected later in the ELISA plate.

Following the below steps we realized a successful solid phase extraction for the oxytocin peptide:
1) Column activation: condition column with 2ml MeOH (each time 1 ml)
2) condition column with 2ml H2O pure (each time 1 ml)

3) Loading samples: load sample up to 1 ml, load slowly!
4) wash with 1ml of 95% $H_2O$ + 5% (of a solution of 25% $NH_4$)
5) wash with 1ml of 95% $H_2O$ + 5% (of a solution of 25% $NH_4$)
6) Elute slowly with 1.75 ml MeOH 100% for capture of Oxytocin (if using a multi-dispenser, elute with 1ml first followed by 0.75 ml).
7) Speedvac at 45°C at 14 bar for 2 hours, check until dryness
8) store the extract at - 80 °C for further ELISA analyses

*Assay procedure*

1) The standards: ST1: 900 ul Assay buffer, ST2 to ST7: 500 ul Assay buffer
2) add 100 ul OCT standard in ST1, vortex for ~1 min then take 500 ul from ST1 to ST2.. repeat the same until ST7 (which will be 1 ml volume after the dilutions)
3) reconstitute the extract with 250 ul of assay buffer, vortex well: approximately for 1 min

General advice:
a) vortex prior to each pipetting
b) while pipetting try to put it in vertical way without touching the well, avoid the walls as the reagents stay sometimes there.
c) wash the tips in the solution you will pipette before pipetting to have a stable amount of the solution
The steps here are from the provided manual by the kit manufacturer: the plate layout sheet (example Table S1) is prepared prior to each assay as a reference to know where to pipette the designated solutions.
1) vortex the tube prior to each pipetting then Pipet 100 µL of Assay Buffer into the NSB and the Bo wells.
2) vortex the tube prior to each pipetting then pipet 100 µL of Standards #1 through #7 into the appropriate wells.
3) vortex the tube prior to each pipetting then Pipet 100 µL of the Samples into the appropriate wells
4) vortex the tube prior to each pipetting then Pipet 50 µL of Assay Buffer into the NSB wells.
5) Pipet 50 µL of the blue Conjugate into each well, except the Blank wells.
6) Pipet 50 µL of the yellow Antibody into each well, except the Blank and NSB wells.
NOTE: Every well used should be Green in color except the NSB wells which should be Blue. The Blank wells are empty at this point and have no color.
7) Shake the plate gently for 15 min in dark (put it back in the sealing bag).
8) Seal the plate and incubate at 4°C for 18-24 hours.
###### 24 Hours max incubation
9) Empty the contents of the wells and wash 400 µL X 3 times. (test first the washing on pilot plate and clean the filling and aspirating tubes if necessary).
10) Add 5 µL of the blue Conjugate to the TA wells.
11) Add 200 µL of the pNpp Substrate solution to every well.
###### Incubate at room temperature for 1 hour without shaking.
12) Add 50 µL of Stop Solution to every well. This stops the reaction and the plate should be read immediately.
13) Read the optical density at 405 nm, preferably with correction between 570 and 590 nm. Blank the empty blank and the NSB wells for reading. They will be subtracted from the optical densities values as a control of the machine reading performances.
14) Use the software for the analyses: use 4 parameters curve for values calculation as the software will estimate automatically the oxytocin concentrations from the standard curve, the optical density OD is an indication of the optical absorbance: more there are oxytocin in the well lower is the OD reading.

15) Correct the final concentrations by the dilution factor. For example: if you elute 1 ml of saliva and you reconstitute by 250 ul assay buffer, then divide the oxytocin concentration by 4 to have the real oxytocin concentration in the 1mL of the saliva sample.

### Supplementary results:

#### *Baseline oxytocin:*

As explained in the main text, we explored whether there was potential confounding covariation between baseline oxytocin levels and the age, gender, group membership, or talking condition of the participants, which may have existed unintentionally. Our analyses showed that no such significant relationship existed between baseline oxytocin levels and age (BLMER: N = 78, estimate = -0.009, $p = 0.377$), gender (BLMER: N = 78, estimate = -1.181, $p = 0.551$), group membership (BLMER: N = 78, estimate = 0.061, $p = 0.794$), or talking condition (BLMER: N = 78, estimate = -0.226, $p = 0.352$). Also, neither the interaction between gender and age nor group and talking was statistically significant (BLMER: N = 78, estimate = 0.006, $p = 0.654$; estimate = -0.047, $p = 0.889$, respectively, Fig. S1).

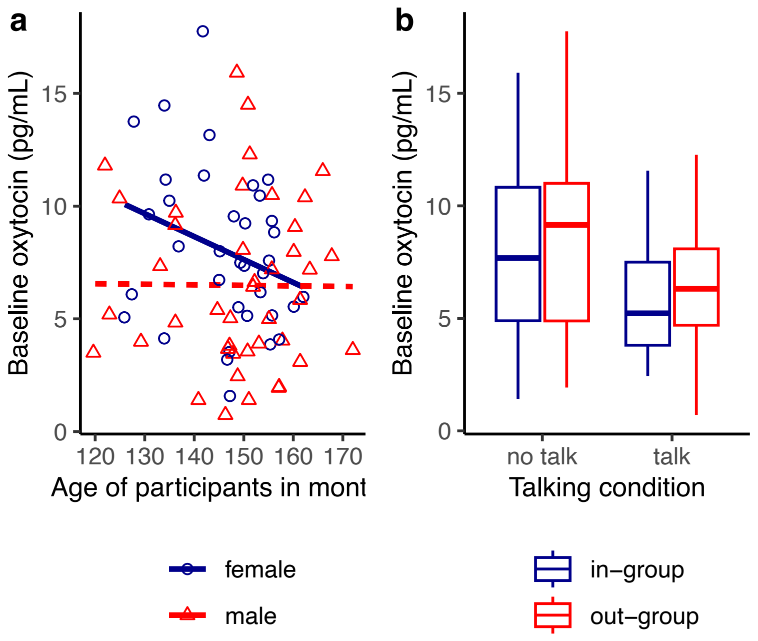


**Figure S1. The relationship between baseline oxytocin levels and potential confounding variables.** (a) scatterplot of baseline oxytocin and age of the participants as a function of gender. (b) boxplot of median and interquartile of baseline oxytocin levels as a function of talking condition and group membership. Statistical analyses shows no significant differences ($p > 0.05$).

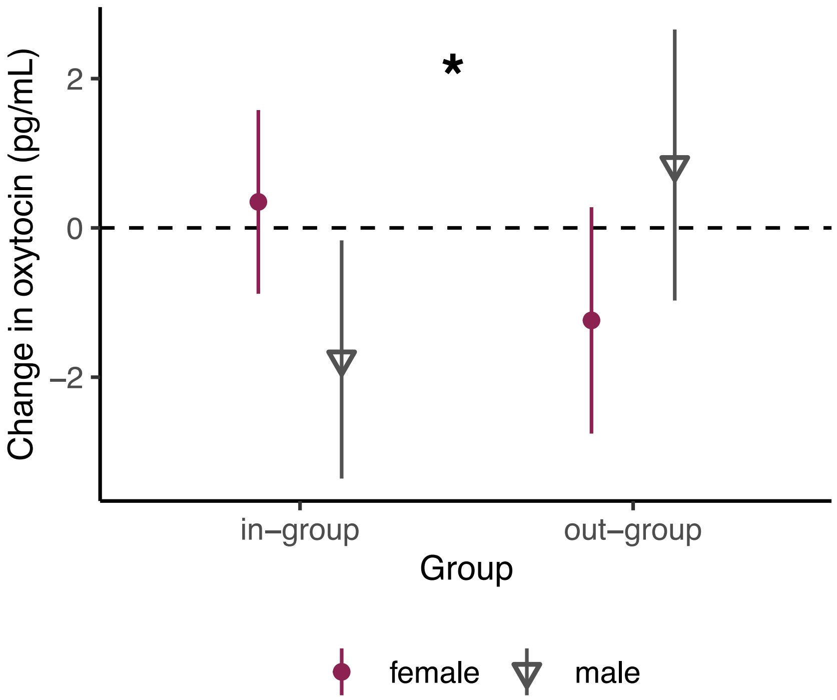


**Figure S2.** Group membership predicting changes in oxytocin levels as a function of participant's gender. Mean and 95% CI of changes in oxytocin from before to after the experiment. The dashed horizontal line refers to zero change in oxytocin levels. *BGLMM $p < 0.05$.

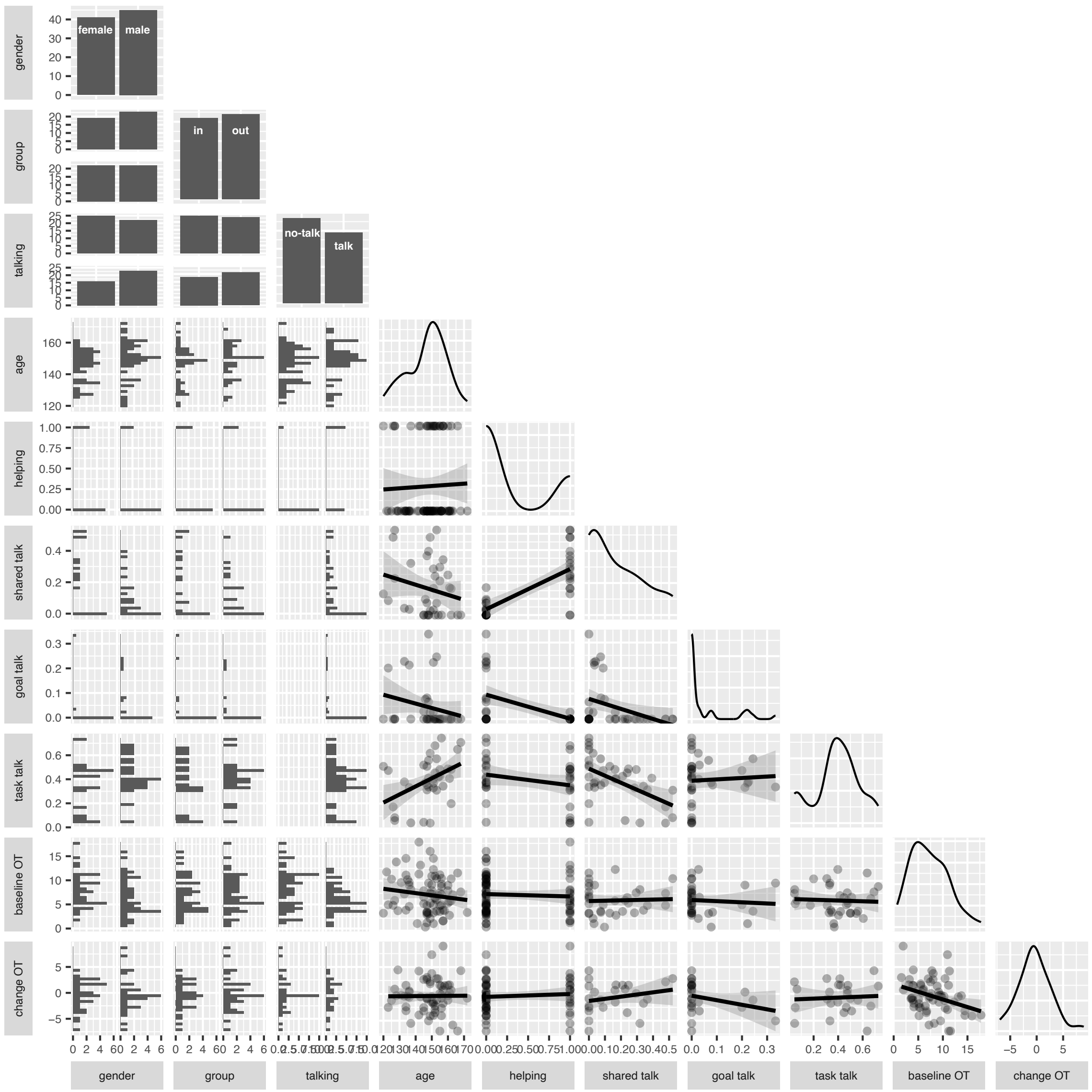


**Figure S3. Correlation plot.**

**Table S1. Summary table of the baseline oxytocin predicting types of talk as a function of group membership and gender.** Significant outcomes are indicated in bold with a threshold of *p < 0.05*. N = 25.

| **bayesian GLMM** | **term** | **estimate** | **lower limit 95 % CI** | **upper limit 95 % CI** | **p-value** |
|---|---|---|---|---|---|
| Shared intentionality talk | **(Intercept)** | **-1.770** | **-3.37** | **-0.18** | **0.029** |
| | Baseline oxytocin | 0.168 | -0.36 | 0.70 | 0.533 |
| | Group (out-group) | -0.750 | -2.42 | 0.92 | 0.380 |
| | Gender (male) | -0.547 | -2.26 | 1.16 | 0.531 |
| | Baseline oxytocin x Group (out-group) | -0.633 | -1.68 | 0.42 | 0.237 |
| | Baseline oxytocin x Gender (male) | 0.279 | -0.77 | 1.33 | 0.601 |
| goal talk | **(Intercept)** | **-9.600** | **-15.60** | **-3.61** | **0.002** |
| | Baseline oxytocin | 4.660 | -2.21 | 11.50 | 0.184 |
| | Group (out-group) | 0.427 | -3.74 | 4.60 | 0.841 |
| | **Gender (male)** | **5.490** | **0.37** | **10.60** | **0.036** |
| | Baseline oxytocin x Group (out-group) | -1.200 | -3.36 | 0.95 | 0.274 |
| | Baseline oxytocin x Gender (male) | -2.440 | -9.18 | 4.30 | 0.478 |
| task talk | **(Intercept)** | **-0.649** | **-1.14** | **-0.16** | **0.010** |
| | Baseline oxytocin | -0.073 | -0.48 | 0.33 | 0.723 |
| | Group (out-group) | 0.217 | -0.31 | 0.74 | 0.417 |
| | Gender (male) | 0.125 | -0.41 | 0.66 | 0.646 |
| | Baseline oxytocin x Group (out-group) | 0.387 | -0.07 | 0.85 | 0.099 |
| | Baseline oxytocin x Gender (male) | -0.284 | -0.74 | 0.17 | 0.223 |

### Supplementary Consent to be signed by the parents (the form is in French).

**FORMULAIRE DE CONSENTEMENT ÉCLAIRÉ POUR L'ÉTUDE**
**"CHASSE AUX ŒUFS DE PÂQUES"**

**Responsables du projet de recherche:**
Prof. Bshary & Prof. Clément
Université de Neuchâtel
Centre de Sciences Cognitives
2000 Neuchâtel

**En quoi consiste cette expérience?**
Cette expérience vise à étudier comment les enfants recherchent de la nourriture et si les niveaux de certaines hormones affectent cette capacité. Nous avons conçu une "Chasse aux Œufs de Pâques" pour les permettre de rechercher des œufs contenant de la "nourriture" imaginaire sous forme de petites vis. Le nombre de vis qu'ils trouveront sera utilisé pour calculer leur récompense. Les consignes de l'expérience leur seront présentées sur l'écran d'un ordinateur et ils pourront poser des questions à tout moment.

**Quelle est la durée de l'expérience?**
L'expérience dure environ 30 minutes.

**Y a-t-il des risques?**
L'expérience consistera à chercher des œufs dans une salle et de fournir deux échantillons de salive. Ces échantillons seront détruits une fois qu'ils auront été analysés. Participer à cette expérience ne comporte donc aucun risque prévisible.

**L'identité de mon enfant est-elle protégée?**
Oui. Leurs données seront maintenues confidentielles lors de l'analyse et de la publication des résultats. Le questionnaire ainsi que la vidéo seront munis d'un numéro anonyme. La vidéo ne sera visionnée que par des personnes impliquées dans ce projet.

Dans certaines situations, il se pourrait que nous montrions un clip de leur vidéo à d'autres chercheurs lors d'une conférence scientifique. Vous n'êtes pas obligés de donner votre accord pour cela, mais si vous l'acceptez, veuillez s'il vous plaît signer ici: _______________________

Au cas où des questions surgiraient après votre participation, veuillez contacter le Dr. Jennifer McClung.

Le responsable de la recherche m'a informé(e) oralement et par écrit des buts de l'étude susnommée, de son déroulement, des effets attendus, des avantages et inconvénients possibles ainsi que des risques éventuels.

J'ai lu et compris les détails au-dessus pour l'étude susnommée. J'ai reçu des réponses satisfaisantes aux questions concernant la participation de mon enfant à cette étude.

J'accepte le fait que les spécialistes responsables travaillant pour promouvoir l'étude, les représentants des autorités et de la commission d'éthique aient un droit de regard sur les données originales concernant mon enfant pour procéder à des vérifications. Ces informations restent toutefois strictement confidentielles.

**FORMULAIRE DE CONSENTEMENT ÉCLAIRÉ POUR L'ÉTUDE**
**"CHASSE AUX ŒUFS DE PÂQUES"**

**Par la présente, je confirme que la participation de mon enfant à cette étude est volontaire. Je dispose du droit d'arrêter l`expérience à tout moment.**

**Expression de consentement**

Je, soussigné(e) …………………………………. (nom) ………………………………. (prénom) consens à ce que mon enfant participe à l'étude "Chasse aux Œufs de Pâques" supervisée par les Prof. Bshary & Clément de l'Université de Neuchâtel.

Neuchâtel, le ......................................................................................................................

Signature du/des parent(s): ...............................................................................................

Je, soussigné(e) …………………………………. (nom) ………………………………. (prénom) consens à ce que mon proche participe à l'étude "Chasse aux Œufs de Pâques" supervisée par les Prof. Bshary & Clément de l'Université de Neuchâtel.

Neuchâtel, le ......................................................................................................................

Signature du responsable légal: .........................................................................................

### Supplementary forms to be filled by the kids for their food preferences.

UNIVERSITÉ DE NEUCHÂTEL

QUESTIONNAIRE

"PRÉFÉRENCES ALIMENTAIRES"

**Entoure ta réponse selon combien tu aimes l'aliment présenté.**

**1. Légumes cuits**

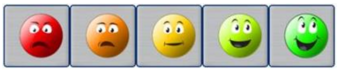

**2. Salades**

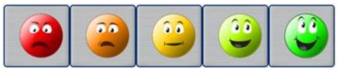

**3. Pâtes**

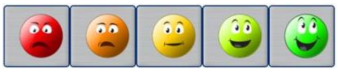

**4. Yoghourts**

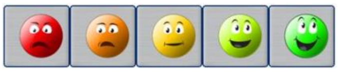

**5. Fruits**

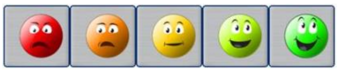

**6. Fromage**

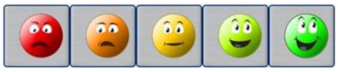

**7. Chocolat**

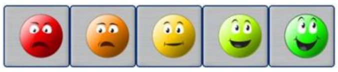

**8. Bonbons**

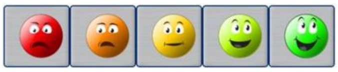

**9. Pain**

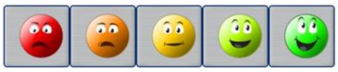

**10. Pizza**

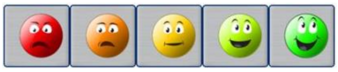
